# Heme uptake in *Lactobacillus sakei* evidenced by a new ECF-like transport system

**DOI:** 10.1101/864751

**Authors:** Emilie Verplaetse, Gwenaëlle André-Leroux, Philippe Duhutrel, Gwendoline Coeuret, Stéphane Chaillou, Christina Nielsen-Leroux, Marie-Christine Champomier-Vergès

## Abstract

*Lactobacillus sakei* is a non-pathogenic lactic acid bacterium and a natural inhabitant of meat ecosystems. Although red meat is a heme-rich environment, *L. sakei* does not need iron or heme for growth, while possessing a heme-dependent catalase. Iron incorporation into *L. sakei* from myoglobin and hemoglobin was formerly shown by microscopy and the *L. sakei* genome reveals a complete equipment for iron and heme transport. Here, we report the characterization of a five-gene cluster (*lsa1836-1840*) encoding a putative metal iron ABC transporter. Interestingly, this cluster, together with a heme dependent catalase gene, is also conserved in other species from the meat ecosystem. Our bioinformatic analyses revealed that the locus might refer to a complete machinery of an Energy Coupling Factor (ECF) transport system. We quantified *in vitro* the intracellular heme in wild-type (WT) and in our Δ*lsa1836-1840* deletion mutant using an intracellular heme sensor and ICP-Mass spectrometry for quantifying incorporated ^57^Fe heme. We showed that in the WT *L. sakei*, heme accumulation occurs fast and massively in the presence of hemin, while the deletion mutant was impaired in heme uptake; this ability was restored by *in trans* complementation. Our results establish the main role of the *L. sakei* Lsa1836-1840 ECF-like system in heme uptake. This research outcome shed new light on other possible functions of ECF-like systems.

**Importance:** *Lactobacillus sakei* is a non-pathogenic bacterial species exhibiting high fitness in heme rich environments such as meat products, although it does not need iron nor heme for growth. Heme capture and utilization capacities are often associated with pathogenic species and are considered as virulence-associated factors in the infected hosts. For these reasons, iron acquisition systems have been deeply studied in such species, while for non-pathogenic bacteria the information is scarce. Genomic data revealed that several putative iron transporters are present in the genome of the lactic acid bacterium *L. sakei.* In this study, we demonstrate that one of them, is an ECF-like ABC transporter with a functional role in heme transport. Such evidence has not yet been brought for an ECF, therefore our study reveals a new class of heme transport system.

## Introduction

Iron is an essential element for almost all living organisms (1) and heme, an iron-containing porphyrin, is both a cofactor of key cellular enzymes and an iron source for bacteria. Many bacteria encode the complete heme biosynthesis pathway to be autonomous for heme production and partly to guarantee their iron supply. However, some others lack heme biosynthetic enzymes and rely on the environment to fulfill their heme requirements. *Lactococcus lactis* and all known *Lactobacilli* are heme-auxotrophic bacteria (2). Also, it is well established that lactic acid bacteria do not require iron to grow (3) and that their growth is unaffected by iron deprivation. Nevertheless, numerous lactic acid bacteria, such as *L. lactis, Lactobacillus plantarum*, or *Enterococcus faecalis*, require exogenous heme to activate respiration growth in the presence of heme (2).

*Lactobacillus sakei* is a non-pathogenic lactic acid bacterium frequently found on fresh meat. *L. sakei* becomes the predominant flora on vacuum packed meat stored at low temperature (4). Interestingly, abundance of *L. sakei* has been shown to prevent growth of undesirable pathogens such as *Listeria monocytogenes* (5, 6), *Escherichia coli* O157:H7 in both cooked and smashed meat (5, 7, 8), and of spoilers such as *Brochothrix thermosfacta* (7, 8). Therefore, this species is often used as a bioprotective culture in meat products. Nevertheless, mechanisms of synergy and competition between species in such complex matrices are still poorly understood (9). Meat, can be considered as a growth medium naturally rich in iron and heme. Quantification of total iron content in raw meat reported a mean of 2.09 mg total iron/100 g for four beef meat cuts in which 87% was heme iron (10). Although *L. sakei* had a tropism for meat and is known to possess a heme-dependent catalase (11), it is considered to be a bacterium that requires neither iron nor heme to grow.

First insights on iron/heme utilization by *L. sakei* came from its whole genome analysis (12) with the identification of coding sequences of several iron transporters, regulators and iron-containing enzymes. Later, microscopy analysis of *L. sakei* cells combined to spectroscopy methods showed that *L. sakei* is able to incorporate iron atoms from complexed iron such as myoglobin, hemoglobin, hematin, and transferrin (13). This suggested that *L. sakei* may display heme or heminic-iron storage ability, although the analytical method used was not quantitative and the precise amount of iron compound that *L. sakei* is able to store was not determined. Hematin did not show any effect on growth of *L. sakei*, but hematin has been shown to prolong bacteria viability in stationary phase (13). However, the mechanisms underlining *L. sakei* survival in the presence of heme need to be unraveled.

Heme acquisition systems have mainly been studied in Gram-negative and Gram-positive pathogens that acquire heme from host hemoproteins in a two steps process (for a review, see (14–16)). First, cell surface or secreted proteins scavenge free heme molecules or complexed heme. Then, transmembrane transporters, generally ATP-binding cassette (ABC) transporters, carry the heme moiety into the intracellular space. Gram-positive bacteria rely mainly on surface-exposed receptors that shuttle heme through the cell-wall and deliver it to an ABC transporter for subsequent transfer into the cytoplasm. Within Gram-positive pathogens, one of the most well characterized heme uptake system is the *Staphylococcus aureus I*ron *S*urface *D*eterminants (Isd) system. The staphylococcal machinery is inserted into a ten-gene locus encoding cell-wall anchored proteins (IsdABCH), a membrane transport system (IsdDEF), a sortase (SrtB) and two cytoplasmic heme-oxygenases (IsdG and IsdI) (17, 18). IsdB and IsdH are responsible for binding host hemoproteins or heme. IsdA extracts heme from IsdB or IsdH and transfers it to IsdC. Funneled heme is finally transferred into the cytoplasm through the membrane by the IsdDEF ABC transporter where it is finally degraded to release free iron by the heme oxygenases IsdG and IsdI. Several of these Isd proteins contain *Nea*r iron *T*ransporter (NEAT) domain, present only in Gram-positive bacteria, and specific to interact with hemoproteins and heme. NEAT domain is a 150-acid residues domain that despite sequence variability displays a conserved β-barrel and a hydrophobic pocket involved in heme binding (19).

Thus far, heme acquisition systems in heme auxotrophic organisms have only been reported for *Streptococci* (15, 20, 21). In *S. pyogenes*, the system involves the Shr and Shp NEAT-domain proteins and the Hts ABC transporter (20, 22, 23). In *Lactococcus lactis*, heme homeostasis, especially heme efflux systems, have been deeply characterized (24, 25). Nevertheless, the acquisition of exogenous heme remains poorly characterized. Heme transport across *L. sakei* membrane is still unknown. Additionally, bioinformatic analysis shows that the genome of *L. sakei* does not contain any NEAT domain (12) which suggests that heme transit could involve transport systems distinct from *Streptococci* and *S. aureus* (14).

Regarding prokaryotic metal ion uptake transporters, comparative and functional genomic analysis have identified Energy-Coupling Factor (ECF) transporters as a novel type of ABC importers widespread in Gram-positive bacteria and first identified in lactic acid bacteria (26). The studies identified genes encoding a ABC-ATPases plus three or four membrane proteins within the same or adjacent to operons, which were implicated in vitamin production or synthesis of metal-containing metalloenzymes (27). Their predicted role in cobalt or nickel ions uptake and delivery within the cell was demonstrated in *Salmonella enterica* and *Rhodoccus capsulatus*, respectively. Since then, ECF-coding genes have been evidenced in *Mycoplasma, Ureaplasma* and *Streptococcus* strains. They were also shown to function as importers not only for transition metal ions but also for vitamins as riboflavin and thiamine (27). Recently, several ECF systems have been characterized, among them folate and pantothenate ECF transport in *Lactobacillus brevis*, and cobalt ECF in *R. capsulatus* (28–31). It was evidenced that ECF transporters constitute a novel family of conserved membrane transporters in prokaryotes, while sharing a similar four domains organization as the ABC transporters. Each ECF displays a pair of cytosolic nucleotide-binding ATPases (the A and A’ components also called EcfA and EcfA’), a membrane-embedded substrate-binding protein (the S component or the EcfS), and a transmembrane energy-coupling component (The T component or EcfT). The quadripartite organization has a 1:1:1:1 stoichiometry. Notably, the S component renders ECF mechanistically distinct from ABC transport systems as it is predicted to shuttle within the membrane, when carrying the bound substrate from the extracellular side into the cytosol (see the recent review (26)). Accordingly, the S-component solely confers substrate specificity to the uptake system (28). Till the 2000s, folate, riboflavin and thiamine ECF importers have been reported for *L. lactis* (32–34). Similarly, folate, hydroxyl pyrimidine and pantothenate ECFs have been reported and structurally characterized for *L. brevis* (28, 30, 31), both Gram-positive rod shape species of lactic acid bacteria. In this paper, we mainly targeted *L. sakei* locus *lsa1836-1840* encoding a putative ABC transporter, and evaluated its role as a heme transport system, combining *in silico* bioinformatics analysis with *in vitro* functional analysis. We showed that this system encodes the complete machinery of an ECF-like importer, including the extracellular proteins that initiate heme scavenging. Furthermore, we were able to dock heme at the binding site formed by the interface of those two extracellular proteins that were homology modeled. In parallel, we quantified the heme storage properties of *L. sakei*, and compared WT *L. sakei* with the Δ*lsa1836-1840 L. sakei* deletion mutant using an intracellular heme-reporter gene and mass-spectrometry quantification of iron-labelled heme. We showed that *L. sakei* Δ*lsa1836-1840* was strongly impaired in its ability to incorporate heme, while its complementation abolished this phenotype. Additionally, when the *lsa1836-1840* genes were overexpressed, heme incorporation was boosted. Thus, we were able to show *in vitro* that this five-gene locus plays an important role in active heme import. To our knowledge, this is the first time that an ECF is reported to being involved in heme incorporation.

## Results

### 1. Putative iron and heme transport systems in *Lactobacillus sakei*

Accurate analysis of the genome of *L. sakei* 23K (12), focused on heme/iron transport systems and heme utilization enzymes, previously led to the identification of 5 putative iron transport systems, five heme transport systems and one heme-degrading enzyme (Table 1). First, two genes *lsa0246* and *lsa1699* encoding proton motive permeases, which belong to the MntH family of manganese uptake, might be involved in iron or heme uptake. Notably, in *L. lactis*, a *mntH* mutant was impaired in Fe^2+^ transport (35).

**Table 1.**

| **Locus tag and Functional category** | **Predicted protein function** |
| --- | --- |
| **ABC transporters** |  |
| *lsa0399-0402* | Fhu |
| *lsa1836-1840* | Putative metal ion ABC transporter, cobalamin transporter |
| *lsa1366-1367* | Putative ABC exporter (heme-efflux machinery) |
| **Proton-motive force transporters** |  |
| *lsa0246* | Mn<sup>2+</sup>/ Zn<sup>2+</sup>/ Fe<sup>2+</sup> transporter |
| *lsa1699* | Mn<sup>2+</sup>/ Zn<sup>2+</sup>/ Fe<sup>2+</sup> transporter |
| **Membrane proteins** |  |
| *lsa1194-1195* | Uncharacterized proteins |
| **Heme-modifying enzyme** |  |
| *lsa1831* | Dyp-type peroxidase |

Second, an operon, composed of the genes *lsa1194-1195* coding for poorly defined membrane proteins of the CCC1 family, is putatively involved in heme or iron transport. In yeast, CCC1 is involved in the manganese and iron ions transport from the cytosol to the vacuole for storage (36). Additionally, the gene *lsa1194*, has been shown to be upregulated in a global transcriptomic analysis of genes differentially expressed in the presence of heme (unpublished data).

Third, two ABC systems homologous to the HrtAB and Pef heme-detoxification systems present in *L. lactis* and *Streptococcus agalactiae* (24, 37) were also identified in *L. sakei* genome. These systems are encoded by the *lsa1366-1367* and *lsa0419-0420* genes, respectively. The sequencing of the *lsa0419-0420* region has confirmed the presence of a frameshift and indicated that these genes are not expressed in *L. sakei* 23K strain. The *lsa1366-1367* gene products are homologous to the *L. lactis Llmg_0625-0624* encoded proteins. The *L. lactis* genes code for the HrtB and HrtA proteins, respectively (24). An *in silico* analysis of Lsa1367 and HrtB indicated that these proteins share 33% of sequence identity and, accordingly, the same fold, as assessed by TOPPRED analysis (38). Particularly, the cytoplasmic-exposed Y168 and Y231 amino-acid residues, shown as important for HrtB-heme interaction in *L. lactis* (25), are also present in Lsa1367, which suggests that these genes might be homologous to the *L. lactis* heme export system.

Last, two iron or heme uptake ABC-transporters were identified. Markedly, the operon *lsa0399-0402* encodes a Fhu system, sharing homology with various orthologous genes and operons encoding complexed iron transport systems, and possibly homologous to the *Listeria monocytogenes* HupCGD system. Also, *L. monocytogenes* shows that HupCGD and Fhu are involved in heme and ferrioxamine uptake, respectively (39).

Then, the ABC system encoded within *lsa1836-1840* genes was automatically annotated as involved in cobalamin transport, whilst it shows some levels of similarities with heme import systems described in Gram-positive bacteria (40–43). At first, we carried out a multiple alignment of all putative substrate-binding lipoproteins encoded in the *L. sakei* 23K genome and noticed that Lsa1839 protein was closely related to Lsa0399 from the Fhu system (data not shown) suggesting a possible link to iron/heme transport. Furthermore, if heme transportation would represent a specific fitness for growth in meat, we wondered whether other meat-borne bacteria would contain a similar cluster in their genome. As shown in Figure 1, comparative genomic analysis revealed that the *lsa1836-1840* genes cluster is present in several species known to harbor a tropism for meat. The most interesting observation is that species harboring the *lsa1836-1840*-like cluster also have in their genome a *katA* gene, encoding a heme dependent catalase, while the other species lacking the cluster, such as *Leuconostoc* and *Lactococcus*, were shown to be deprived of catalase-encoding gene. Although such co-occurrence could not constitute a proof of the role of the *lsa1836-1840* cluster in heme transport, this analysis provided an additional argument consolidating this hypothesis.

**Figure 1:**
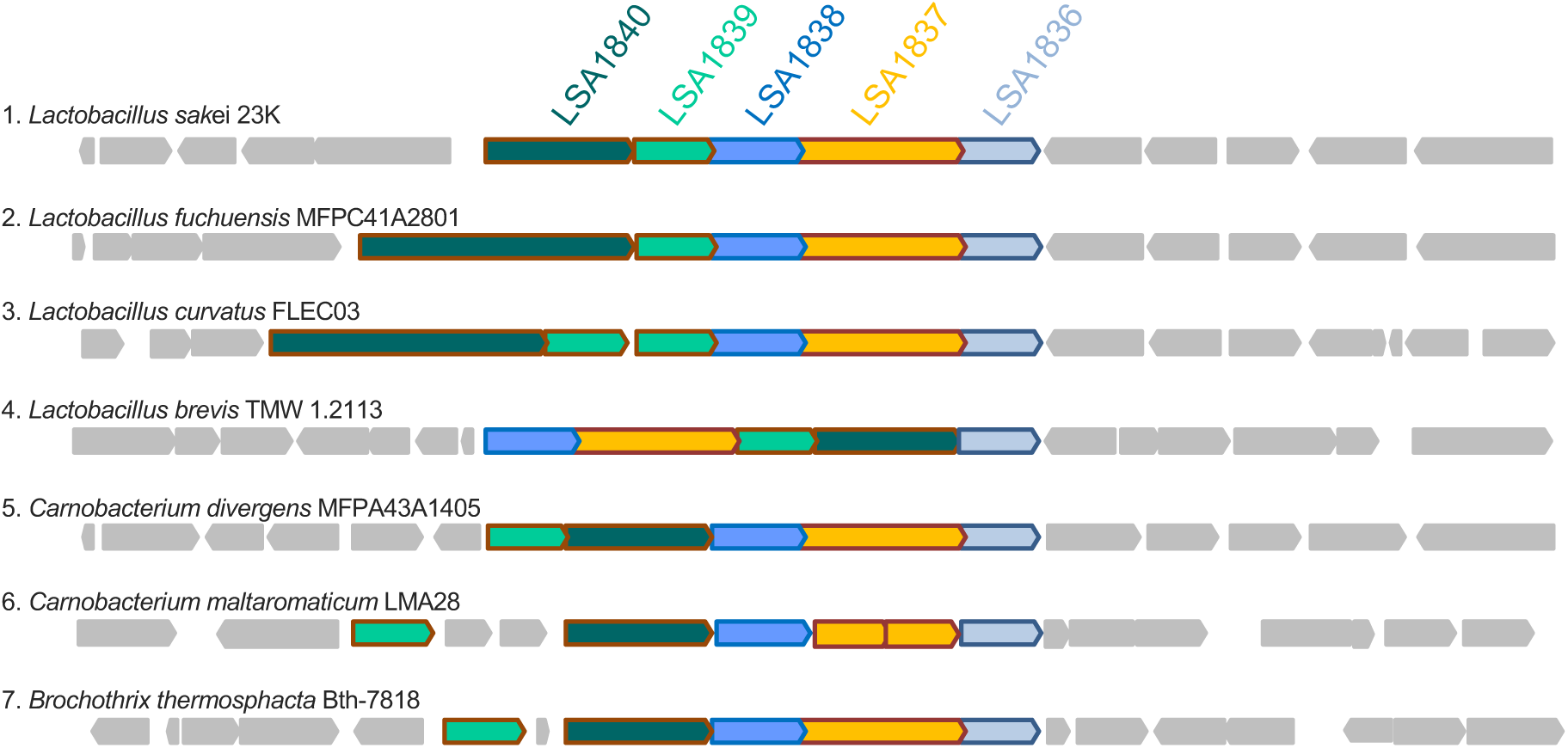
Gene synteny within and around the *lsa1836-1840* gene cluster of *L. sakei* 23K with other Gram-positive species found frequently on meat products. Genes in grey background are unrelated to this cluster and are not conserved between the different genomes. The name of the species and of the strains used for analysis are depicted on the right. All of this species contains a *katA* gene (encoding a heme-dependent catalase) in their genome. Other meat-borne species including *Leuconostoc, Lactococcus, Vagococcus* species also found on meat are not shown due to the lack of both *katA* gene and *lsa1836-1840* gene cluster.

### 2. The *lsa1836-1840* encodes an ECF like transport system putatively involved in heme transport

Due to the conservation of the operon *lsa1836-1840*, each of the five sequences was analyzed comprehensively using bioinformatics. It includes multiple sequence alignment, as well as 3D structure, proteins network and export peptide predictions. Lsa1836 shows a sequence similarity of more than 30%, associated to a probability above 99% with an e-value of 8. e^-15^, to share structural homology with the membrane-embedded substrate-binding protein component S from an ECF transporter of the closely related *L. brevis*, as computed by HHpred (44). Accordingly, its sequence is predicted to be an integral membrane component with six transmembrane helices, and a very high rate of hydrophobic and apolar residues, notably 11 tryptophan amino-acid residues among the 230 residues of the full-length protein (Fig. 2A). HHpred analysis indicates that Lsa1837 shares more than 50 % sequence similarity with the ATPase subunits A and A’ of the same ECF in *L. brevis* (Fig. 2A). With 100% of probability and a e-value of 1. e^-35^, Lsa1837 describes two repetitive domains, positioned at 9-247 and 299-531, where each refers structurally to one ATPase very close in topology to the solved ATPase subunits, A and A’ of ECF from *L. brevis*, respectively. Appropriately, the N-terminal and C-terminal ATPases, are predicted to contain an ATP-binding site. Lsa1837 could correspond to the fusion of ATPase subunits, A and A’. Protein Lsa1838 shows sequence similarity of above 30%, with a probability of 100 % and e-value of 1. e^-30^, to share structural homology with the membrane-embedded substrate-binding protein component T from the ECF transporter of *L. brevis* (Fig. 2A). Interestingly, similar bioinformatic analysis of sequence and structure prediction demonstrates that Lsa1839 and Lsa1840 share both 99.8% structural homology, and e-value of 1. e^-24^ and of 1. e^-21^, with the β and α domains of human transcobalamin, respectively (Fig. 2A). Consistently, both proteins have an export signal located at their N-terminal end. Taken together, these results predict with high confidence that the transcriptional unit encodes the complete machinery of an ECF, including the extracellular proteins that initiate the scavenging of iron-containing heme (Fig. 2A). Each protein compartment is predicted through the presence/absence of its signal peptide as being extracellular, embedded in the membrane or cytosolic. Correspondingly, every protein sequence associates appropriate subcellular location with predicted function. In line with that, the network computed by String for the set of proteins of the operon shows that they interact together from a central connection related to Lsa1837, which corresponds to the ATP-motor couple of ATPases (45). The transcriptional unit also encompasses Lsa1839 and Lsa1840, highly homologous to β and α subunits of transcobalamin respectively, that are highly hypothesized to initiate the scavenging of heme from the extracellular medium. To address the capacity of those subunits of transcobalamin-like binding domain to bind a heme moiety, we homology-modeled Lsa1839 and Lsa1840. We then assembled the biological unit composed of the heterodimer formed by β and α subunits, using the related 3D templates of corresponding subunit of haptocorrin and transcobalamin. Subsequently, an iron-containing heme moiety was docked into the groove, located at the interface of the complex formed by the two proteins. The redocking of cobalamin in haptocorrin and cyanocobalamin in transcobalamin shows a binding energy of -17 and -12 kcal/mol, respectively (Fig. 2B). With a binding energy of -9 kcal/mol, the heme bound to the crevice formed by Lsa1839 and Lsa1840 displays an affinity in the same range than the endogenous ligands, and emphasizes that the assembly composed of Lsa1839 and Lsa1840 could be compatible with the recognition and binding of a heme (Fig. 2B). To resume, Lsa1836-1840 describes a complete machinery that could be able to internalize a heme instead or additionally to a cobalamin molecule. Importantly, this operon includes also the extracellular scavenging β- and α-like subunits of transcobalamin, which advocates for that the S-component Lsa1836 is possibly very specific for iron-containing heme. In line with that, despite a closely conserved fold, the S-component does not display the strictly conserved residues known to bind cobalt-containing cobalamin.

**Figure 2:**
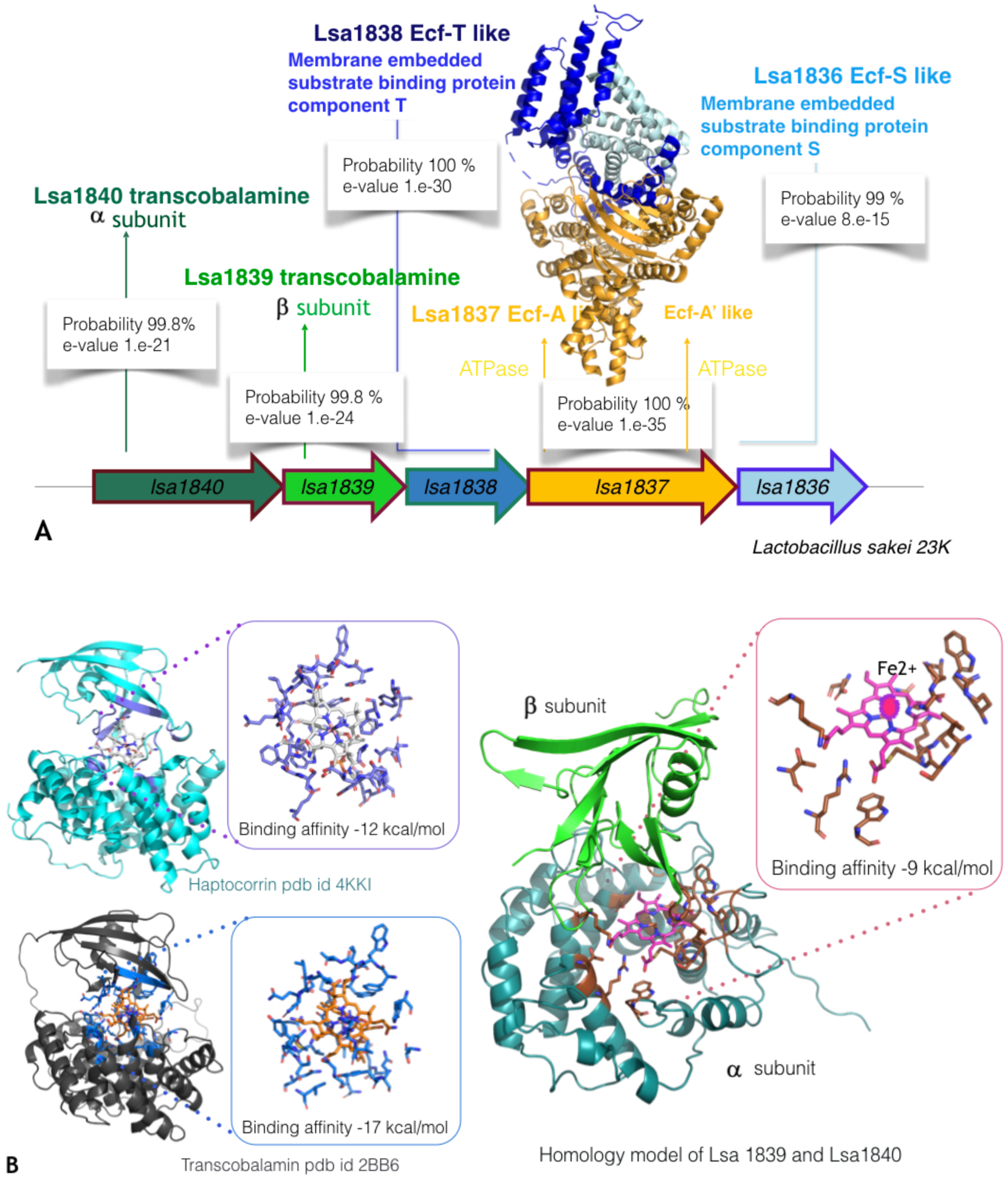
Panel A details the structural and functional bioinformatic assessment for each gene of the *lsa1836-1840* operon. Panel B focuses on Lsa1839 and Lsa1840 and highlights (left) the binding interaction and affinity of the human haptocorrin with cyano-cobalamin and bovine transcobalamin with cobalamin, respectively. They were used as 3D template and positive control for the modeling of transcobalamin-like proteins Lsa1840 and Lsa1839. Panel B (right) shows the best pose of iron containing heme as computed by Autodock4 within the binding pocket formed at the interface of a and b subunits of homology modeled Lsa1840 and Lsa1839, respectively.

No heme synthesis enzymes are present in *L. sakei* genome, nevertheless a gene coding for a putative heme-degrading enzyme of the Dyp-type peroxidase family, *lsa1831*, was identified in the *L. sakei* genome. Its structure is predicted to be close to DypB from *Rhodococcus jostii* (46). Interestingly, residues of DypB involved in the porphyrin-binding, namely Asp153, His226 and Asn246, are strictly conserved in Lsa1831 (47). Markedly, the *lsa1831* gene is located upstream of the *lsa1836*-*1840* operon putatively involved in the active heme transport across the membrane.

Our bioinformatical analysis allows the functional reannotation of the *lsa1836-1840* genes into the complete machinery of an Energy-Coupling Factor, possibly dedicated to the transport of iron through the heme (Fig. 3A-B). Consistently, the Lsa1831 enzyme, which is close to the *lsa1836-1840* loci, could participate downstream to release iron from the heme once inside the cytoplasm.

**Figure 3:**
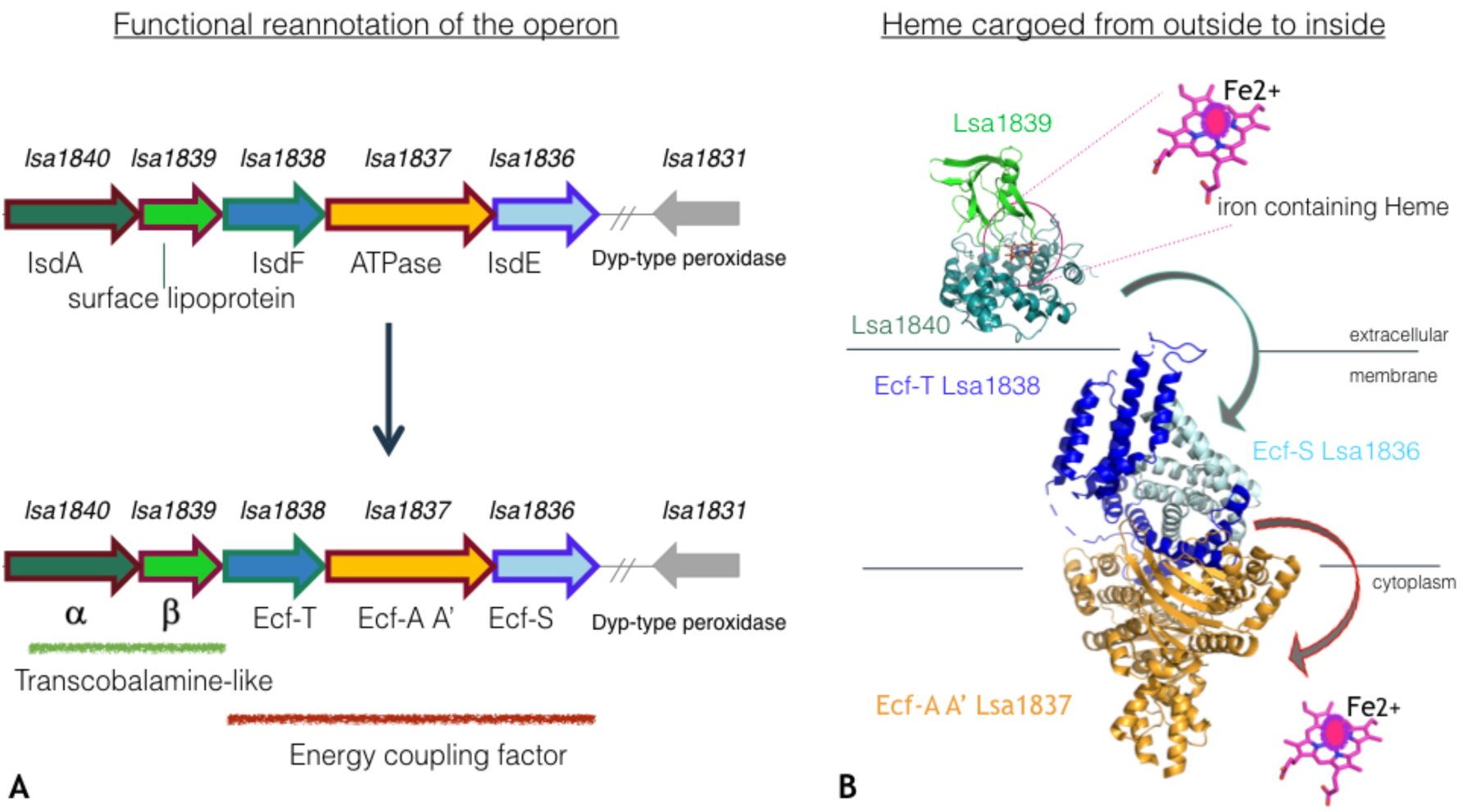
A, Functional reannotation of the operon *lsa1836-1840* from *L. sakei* 23K after serial analysis of 3D structure/function prediction for each gene of the operon. B, Reconstitution of iron-containing heme transport, initially scavenged between the a and b subunits of the transcobalamine-like transporter, coded by *lsa1839-1840*, then cargoed from the extracellular into the intracellular compartments through the complete ECF machinery coded by *lsa1836-1838* portion of the operon. Possibly, gene *lsa1831* positioned in the vicinity of the loci *lsa1836-1840* could code for a protein Dyp-type peroxidase that ultimately releases the iron from the heme.

### 3. The Lsa1836-1840 is *in vitro* an effective actor of heme uptake in *L. sakei*

To confirm the above transporter as involved in heme trafficking across the membrane, a *lsa1836-1840* deletion mutant was constructed by homologous recombination. The *L. sakei* Δ*lsa1836-1840* mutant was analyzed for its capacity to internalize heme using an intracellular heme sensor developed by Lechardeur and co-workers (24). This molecular tool consists in a multicopy plasmid harboring a transcriptional fusion between the heme-inducible promoter of *hrtR*, the *hrtR* coding sequence and the *lacZ* reporter gene, the pP_hrt_ *hrtR-lac* (Table 2). In *L. lactis*, HrtR is a transcriptional regulator that represses the expression of a heme export system, HrtA and HrtB, as well as its own expression in the absence of heme. Upon heme binding, the repression is alleviated allowing the expression of the export proteins (24). As *L. sakei* possesses the *lacLM* genes, it was necessary to construct the Δ*lsa1836-1840* mutant in the *L. sakei* RV2002 strain, a *L. sakei* 23K Δ*lacLM* derivative, yielding the RV4057 strain (Table 2). The pP_hrt_ *hrtR-lac* was then introduced in the RV2002 and RV4057 strains, yielding the RV2002 *hrtR-lac* and the RV4057 *hrtR-lac* strains (Table 2). β-Galactosidase (β-Gal) activity of the RV4057 *hrtR-lac* strain grown in a chemically defined medium (MCD) (48) in the presence of 0.5, 1 and 5 µM hemin was determined and compared to that of the RV2002 *hrtR-lac* used as control (Fig. 4A). We showed that hemin reached the intracellular compartment as β-Gal expression was induced by hemin. Relative β-Gal activity of the RV4057 *hrtR-lac* mutant strain showed a slight increase as compared to the WT at 0.5 µM heme but a statistically significant two-fold reduction was measured at 1 µM heme and further, a 40% reduced activity was shown at higher hemin concentration. This indicates that the intracellular abundance of heme is significantly reduced in the RV4057 bacterial cells at 1 and 5 µM heme, while it is similar to the WT at low heme concentrations. The method described above did not allow to quantify the absolute amount of heme incorporated by bacteria as only cytosolic heme may interact with HrtR. Therefore, we used hemin labeled with the rare ^57^iron isotope (^57^Fe-Hemin) combined with Inductively Coupled Plasma Mass Spectrometry (ICP-MS) to measure with accuracy the total heminic-iron content of cells. Quantification of ^57^Fe was used as a proxy to quantify heme. The absolute number of heme molecules incorporated by the Δ*lsa1836-1840* mutant was also quantified using ^57^Fe-hemin. The Δ*lsa1836*-*1840* mutant was constructed in the WT *L. sakei* 23K genetic background to obtain the RV4056 strain (Table 2). Bacteria were incubated in the MCD, in the absence or in the presence of 1, 5 or 40 µM of ^57^Fe-hemin. ICP-MS quantification indicated that the ^57^Fe content of the two strains was similar at 1 µM ^57^Fe-hemin. A 5-fold reduction in the ^57^Fe content of the RV4056 strain was measured at 5 µM heme concentrations and a 8-fold at 40 µM heme, by comparison with the WT (Fig. 4B).

**Table 2:**
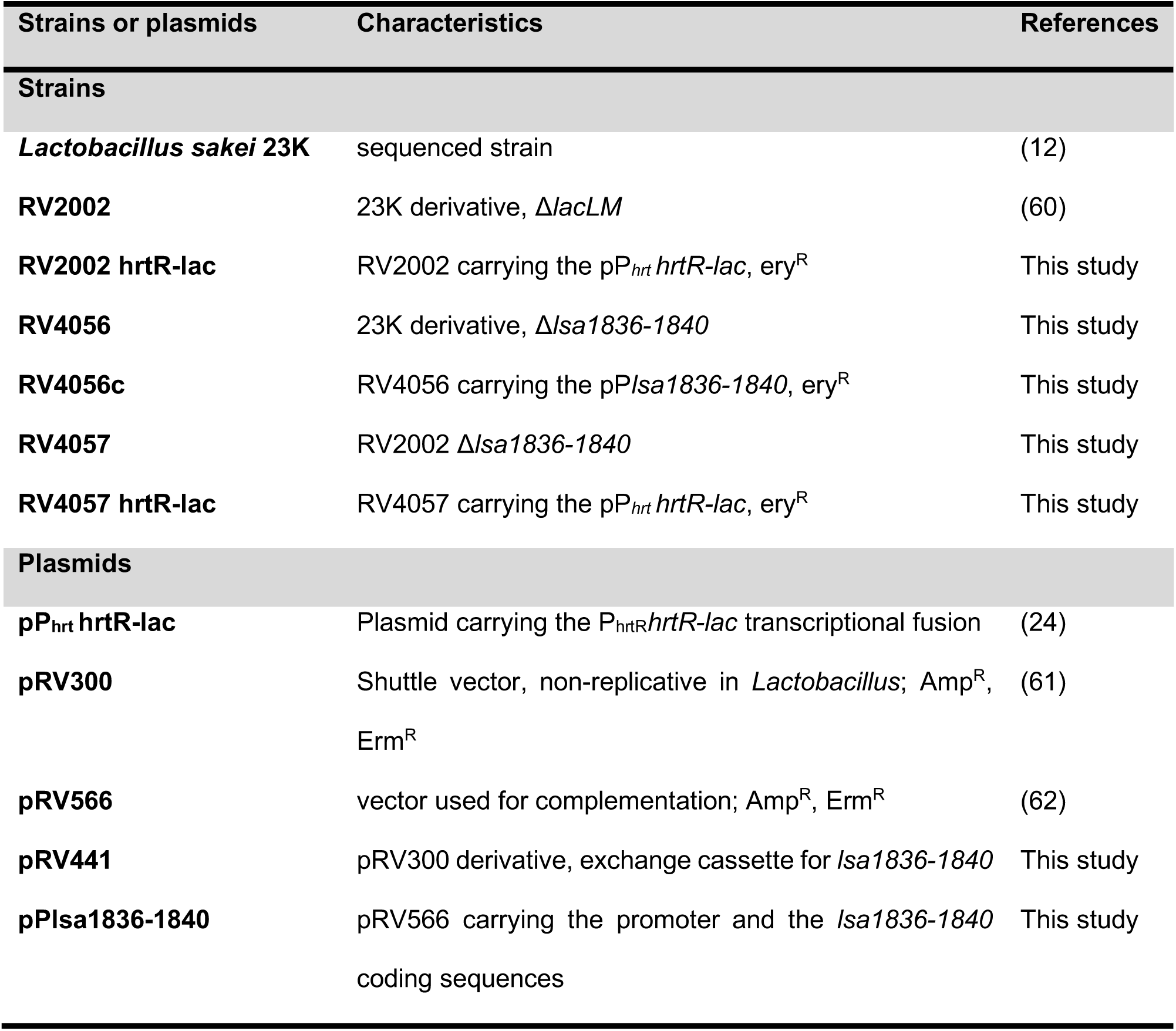
Strains and plasmids used in this study

**Figure 4:**
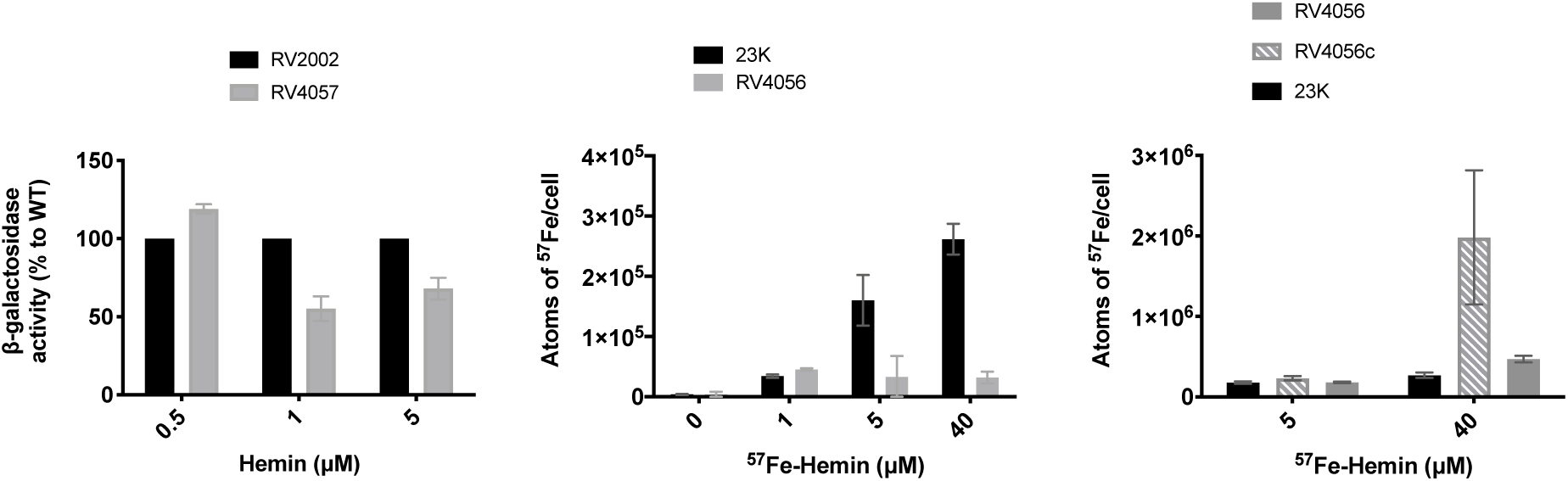
Heme incorporation is reduced in the Δ*lsa1836-1840 L. sakei* deletion mutant. A, *In vivo* detection of intracellular heme content of the RV2002 and Δ*lsa1836-1840* (RV4057) mutant strains. Strains carrying the pP_*hrtR*_*hrtR-lac* were grown in hemin and β-Gal activity was quantified by luminescence (see “Material and methods”). For each experiment, values of luminescence obtained with no added hemin are subtracted and β-Gal activity of strains was expressed as the percentage to the RV2002 strain for each hemin concentration. Mean values are shown (n=3). Error bars represent the standard deviation. B, Quantification of the ^57^Fe content of the WT (23K) and the Δ*lsa1836-1840* (RV4056) strains grown in the absence and presence of indicated ^57^Fe-hemin concentrations. Results represent the mean and range from at least two independent experiments. C, Quantification of the ^57^Fe content of the WT (23K), the Δ*lsa1836-1840* (RV4056) and the Δ*lsa1836-1840* pP*lsa1836-1840* (RV4056c) strains grown in the absence and presence of indicated ^57^Fe-hemin concentrations. Results represent the mean and range of two independent experiments.

To confirm the major role of the *lsa1836-1840* gene products in heme acquisition, we analyzed the ^57^Fe content of the RV4056 strain harboring the pP*lsa1836-1840*, a multicopy plasmid that expresses the *lsa1836-1840* operon under its own promoter, and compared it to the WT. The quantification of the ^57^Fe atoms in the RV4056 pP*lsa1836-1840* bacteria shows a 1.3 time and a 7 times higher iron content at 5 and 40 µM ^57^Fe-hemin, respectively, by comparison with measurements done on WT bacteria (Fig. 4C).

These experiments confirm that the Lsa1836-1840 system is involved *in vitro* in the active incorporation of heme in *L. sakei*.

### 4. Heme accumulates inside the *L. sakei* cytosol at low heme concentrations

We then addressed the ability for *L. sakei* to consume heme or iron to survive. We knew from a previous study that *L. sakei* incorporates preferentially heminic-compounds from the medium, probably as an adaptation to its meat environment (13). Data obtained previously showed that the incorporation of heme molecules are qualitatively correlated with both the concentration of heme in the growth medium, and the survival properties of the bacteria in stationary phase, suggesting that *L. sakei* could use heme or iron for its survival (See Supplemental text, Fig. S1 and S2). Nevertheless, heme incorporation could not be quantified with accuracy in the previous studies. To tackle that, the intracellular heme levels incorporated by *L. sakei* were quantified. The RV2002 *hrtR-lac* strain (Table 2) was grown in MCD in the presence of increasing concentration of hemin, and the β-Gal activity of cells was measured (Fig. 5A). We showed that the β-Gal activity increased with the concentration of the hemin molecule in the growth medium. A plateau was reached when cells were grown in 0.75 - 2.5 µM hemin. Incubation of cells in higher hemin concentrations did not allow to increase further β-Gal activity.

**Figure 5:**
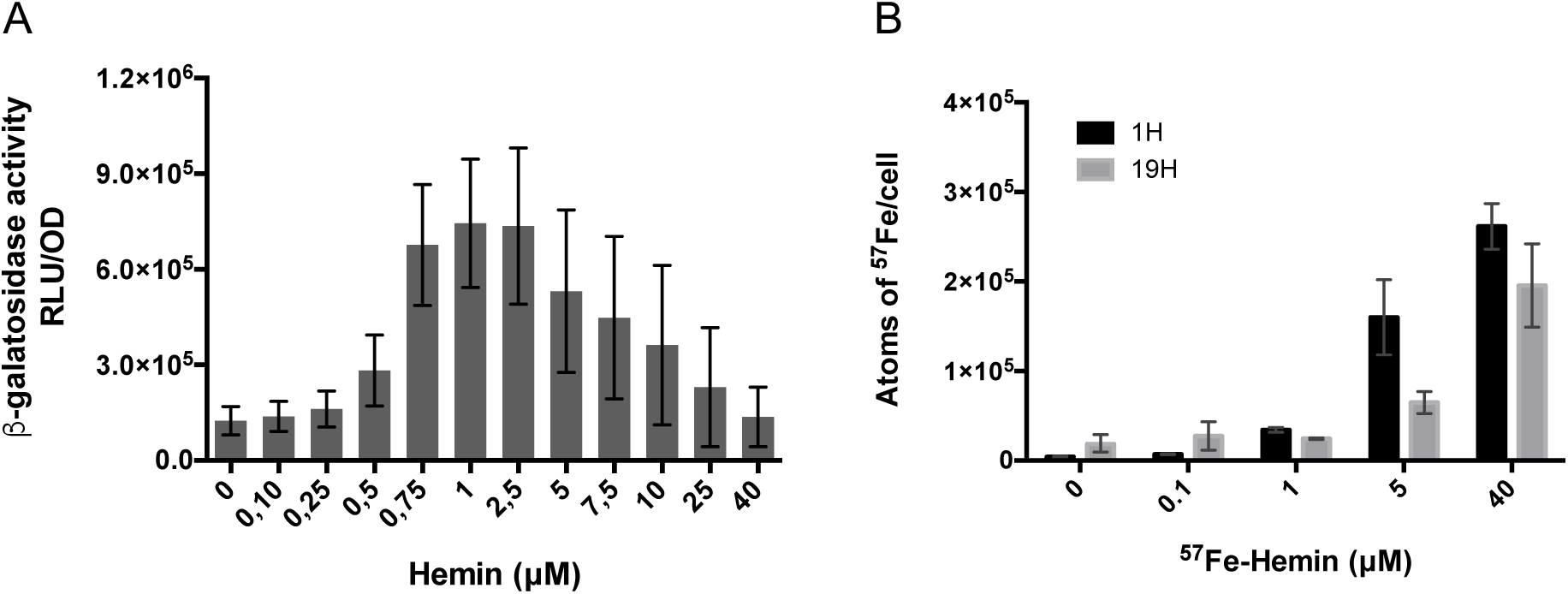
Quantification of heme incorporation in *L. sakei*. A, *In vivo* detection of intracellular hemin molecules through the expression of the *lacZ* gene. The *L. sakei* RV2002 hrtR-lac strain was grown for 1 h in the presence of the indicated concentrations of hemin. β-Gal activity was quantified by luminescence (see “Material and methods”). Mean values are shown (n=7). Error bars represent the standard deviation. B, Quantification of the ^57^Fe content of the WT (23K) strain grown in the absence and presence of ^57^Fe-hemin for 1h and 19h. The mean values and range of two independent experiments are shown. RLU, relative light units.

### 5. Heme incorporation in *L. sakei* is rapid and massive

The absolute number of heme molecules incorporated by *L. sakei* 23K (Table 2) was also quantified using ^57^Fe-hemin. Cells were grown in MCD in the presence of labeled-hemin. Measurements of the ^57^Fe content of cells showed that the incorporation of ^57^Fe-Hemin is massive and rapid as bacteria are able to incorporate about 35,000 ^57^Fe atoms of heminic origin, within 1 hour in the presence of 1 µM ^57^Fe-Hemin (Fig. 5B). The iron content of cells increased to 160,000 and 260,000 atoms in average when bacteria were grown in a medium containing 5 and 40 µM of ^57^Fe-Hemin, respectively. This indicates that the ^57^Fe content of *L. sakei* cells increased with the ^57^Fe-Hemin concentration in the medium on the 1 to 40 µM range. Measurements of the iron content of bacteria growing in presence of ^57^Fe-Hemin for an extended period of time (19h) did not show additional ^57^Fe accumulation in the bacteria (Fig. 5B). Instead, the number of ^57^Fe atoms associated with bacteria decreased over time highlighting the fact that a massive incorporation of labeled-hemin occurs rapidly after bacteria being in contact with the molecules.

## Discussion

Heme acquisition systems are poorly documented in lactic acid bacteria, probably because heme or iron are not mandatory for growth of these bacterial species, at least under non-aerobic conditions. However, acquisition of exogenous heme allows numerous lactic acid bacteria, among them *L. lactis* and *Lactobacillus plantarum*, to activate, if needed, a respiratory metabolism, when grown in the presence of oxygen (2, 49, 50). This implies that heme has to cross the thick cell-wall of these Gram-positive organisms and may require heme transporters. Thus far, heme acquisition systems in heme auxotrophic organisms have only been reported for *Streptococci* (20, 21) and *S. pyogenes*, where they both involve Shr and Shp NEAT-domain proteins and Hts ABC transporter (20, 22, 23). In lactic acid bacteria, no such functional heme transport has been identified so far. NEAT domains have been identified in several species of lactic acid bacteria, including 15 *Lactobacillus*, 4 *Leuconostoc* and one *Carnobacterium* species, but our study confirmed that *L. sakei* proteins are devoid of such domains (19).

In *L. lactis*, the *fhuCBGDR* operon has been reported to be involved in heme uptake as a *fhuD* mutant is defective in respiration metabolism, suggesting a defect in heme import (15). A genome analysis of several lactic acid bacteria has revealed that a HupC/FepC heme uptake protein is present in *L. lactis, L. plantarum, Lactobacillus brevis* and *L. sakei* (15). This latter in *L. sakei* 23K may correspond to locus *lsa0399* included in a *fhu* operon. An IsdE homolog has also been reported in *L. brevis* genome but the identity of this protein has not been experimentally verified (15).

The genome analysis of *L. sakei* 23K (12), when focused on heme/iron transport systems and heme utilization enzymes, led to the identification of several putative iron transport systems, heme transport systems and heme-degrading enzymes. This heme uptake potential is completely consistent within the meat environment-adapted *L. sakei*. Similarly, the membrane transport system encoded by the *lsa1194-1195* genes, whose function is poorly defined, seems to be important for the bacterial physiology as a *lsa1194-1195* deletion affects the survival properties of this strain (see Supplemental text, Fig. S3 and Fig. S4). Meanwhile, here, we report that the transcriptional unit *lsa1836-1840* shows exquisite structure/function homology with the cobalamin ECF transporter, a new class of ATP-binding cassette importer recently identified in the internalization of cobalt and nickel ions (Fig. 2 and Fig. 3). Indeed, a comprehensive bioinformatics analysis indicates/supports that the *lsa1836-1840* locus codes for 5 proteins that assemble together to describe a complete importer machinery called Energy Coupling Factor. Any canonical ECF transporter comprises an energy-coupling module consisting of a transmembrane T protein (EcfT), two nucleotide-binding proteins (EcfA and EcfA’), and another transmembrane substrate-specific binding S protein (Ecsf). Indeed, Lsa1836-Lsa1838 shows high structural homology with Ecf-S, EcfA-A’ and Ecf-T, respectively. Despite sharing similarities with ABC-transporters, ECF transporters have different organizational and functional properties. The lack of soluble-binding proteins in ECF transporters differentiates them clearly from the canonical ABC-importers. Nevertheless here, *lsa1839* and *lsa1840* code for proteins structurally close to β and α subunits of transcobalamin-binding domain, respectively. They are highly suspected to be soluble proteins dedicated to scavenge heme from the extracellular compartment and we hypothesize that they could bind it and then transfer it to Ecf-S component coded by *lsa1836* (Fig. 3). In line with that, the heterodimer composed of Lsa1839 & Lsa1840, possibly β and α subunits, respectively, have been modeled *in silico* and were shown to accommodate with high affinity an iron-heme ligand at the binding site located at the interface of the two proteins.

Internalization of the cobalt and nickel divalent cations through porphyrin moiety *via* this new class of importer has been demonstrated in lactic acid bacteria, such as *L. lactis* and *L. brevis.* However, nothing was known for the internalization/incorporation of iron-containing heme. A functional analysis of the *lsa1836-1840* gene products was undertaken using Δ*lsa1836-1840* deletion mutant and a complemented strain. Our experiments indicate that the intracellular abundance of heme is significantly reduced in Δ*lsa1836-1840* mutant bacterial cells at 1 and 5 µM heme, while it is similar to the WT at low heme concentrations. Reversely, the mutant strain in which *lsa1836-1840* is expressed from a multicopy plasmid, showed an increase in the heme uptake. Taken together, these experiments confirm that the Lsa1836-1840 system is involved *in vitro* in the active incorporation of heme in *L. sakei*. Also, our syntheny analysis for this operon shows that this feature could be shared within several Gram-positive meat-borne bacteria.

Additionally, we were able to quantify the amount of heme internalized in the three genetic contexts using isotope-labeled hemin and ICP-MS as well as to evaluate the intracellular content of heme using the transcriptional fusion tool. We observed that the intracellular abundance of heme increases with the concentration of heme in the growth medium and can be detected with the intracellular sensor in the 0 - 2.5 µM heme range (Fig. 5A). The drop in the β-gal activity at higher heme concentrations may result from regulation of heme/iron homeostasis either through exportation of heme, degradation of the intracellular heme or storage of the heme molecules, making them unable to interact with HrtR and promoting *lacZ* repression. However, data obtained with the intracellular sensor at higher heme concentration (5-40 µM) contrast with microscopic observations (Fig. S2) and ICP-MS measurements (Fig. 5B) that reported a higher heminic-iron content in cells grown in 40 µM heme than in 5 µM. Indeed, β-gal activity reflecting the abundance of intracellular heme was maximal when cells were grown in a medium containing 1-2.5 µM hemin (Fig. 5A), while ICP-MS measurements showed a 4.5 fold and 8 fold higher number of ^57^Fe atoms in bacteria growing in 5 µM or 40 µM ^57^Fe-Hemin, respectively, than in 1 µM ^57^Fe-Hemin (Fig. 4B). These data are in good agreement with EELS analysis (Fig. S2), which strengthens the hypothesis that heme homeostasis occurs in *L. sakei* and that the incorporated heme molecules would be degraded while iron is stored inside iron storage proteins like Dps, of which orthologous genes exist in *sakei.* Thus iron is detected in *L. sakei* cells but not bound to heme and unable to interact with the intracellular heme sensor HrtR. Storage of heme inside membrane proteins is still an open question as *L. sakei* does not contain cytochromes nor menaquinones (12).

Further analysis is required not only to decipher the exact role of these proteins during the different steps of heme transport across the *L. sakei* membrane and the fate of heme inside *L. sakei* cells, but also to understand the molecular specificity of the Lsa1836-1840 machinery towards iron-containing heme *versus* cobalamin.

## Materials and methods

### Bacterial strains and general growth conditions

The different bacterial strains used throughout this study are described in Table 1. *Lactobacillus sakei* and its derivatives (RV2002 RV2002 hrtR-lac RV4056 RV4056c RV4057 RV4057 hrtR-lac) were propagated on MRS (2) at 30°C. For physiological studies the chemically defined medium MCD (3) supplemented with 0.5% (wt/vol) glucose was used. MCD contains no iron sources but contains possible traces of iron coming from various components or distilled water. Incubation was performed at 30°C without stirring. Cell growth and viability of cells in stationary phase were followed by measuring the optical density at 600 nm (OD_600_) on a visible spectrophotometer (Secoman) and by the determination of the number of CFU ml^-1^ after plating serial dilutions of samples on MRS agar. When needed, media were supplemented with filtered hemin or hematin (Sigma-Aldrich) or with ^57^Fe-hemin (Frontier Scientific) solutions resuspended in 50 mM NaOH. *Escherichia coli* K-12 strain DH5α was used as the host for plasmid construction and cloning experiments. *E. coli* cells were chemically transformed as previously described (4). *L. sakei* cells were transformed by electroporation as previously described (5). For routine growth, *E. coli* strain was propagated in LB at 37°C under vigorous shaking (175 rpm). The following concentrations of antibiotic were used for bacterial selection: kanamycin at 20 µg/mL and ampicillin at 100 µg/mL for *E. coli* and erythromycin at 5 µg/mL for *L. sakei.*

### DNA manipulations

Chromosomal DNA was extracted from Ls cells with DNA Isolation Kit for Cells and Tissues (Roche, France). Plasmid DNA was extracted from *E. coli* by a standard alkaline lysis procedure with NucleoSpin^®^ Plasmid Kit (Macherey Nagel, France). PCR-amplified fragments and digested fragments separated on 0.8% agarose gels were purified with kits from Qiagen (France). Restriction enzymes, *Taq* or *Phusion* high-fidelity polymerase (ThermoScientific, France) and T4 DNA ligase (Roche) were used in accordance with the manufacturer’s recommendations. Oligonucleotides (Table 3) were synthesized by Eurogentec (Belgium). PCRs were performed in Applied Biosystems 2720 Thermak thermocycler (ABI). Nucleotide sequences of all constructs were determined by MWG - Eurofins (Germany).

**Table 3:**
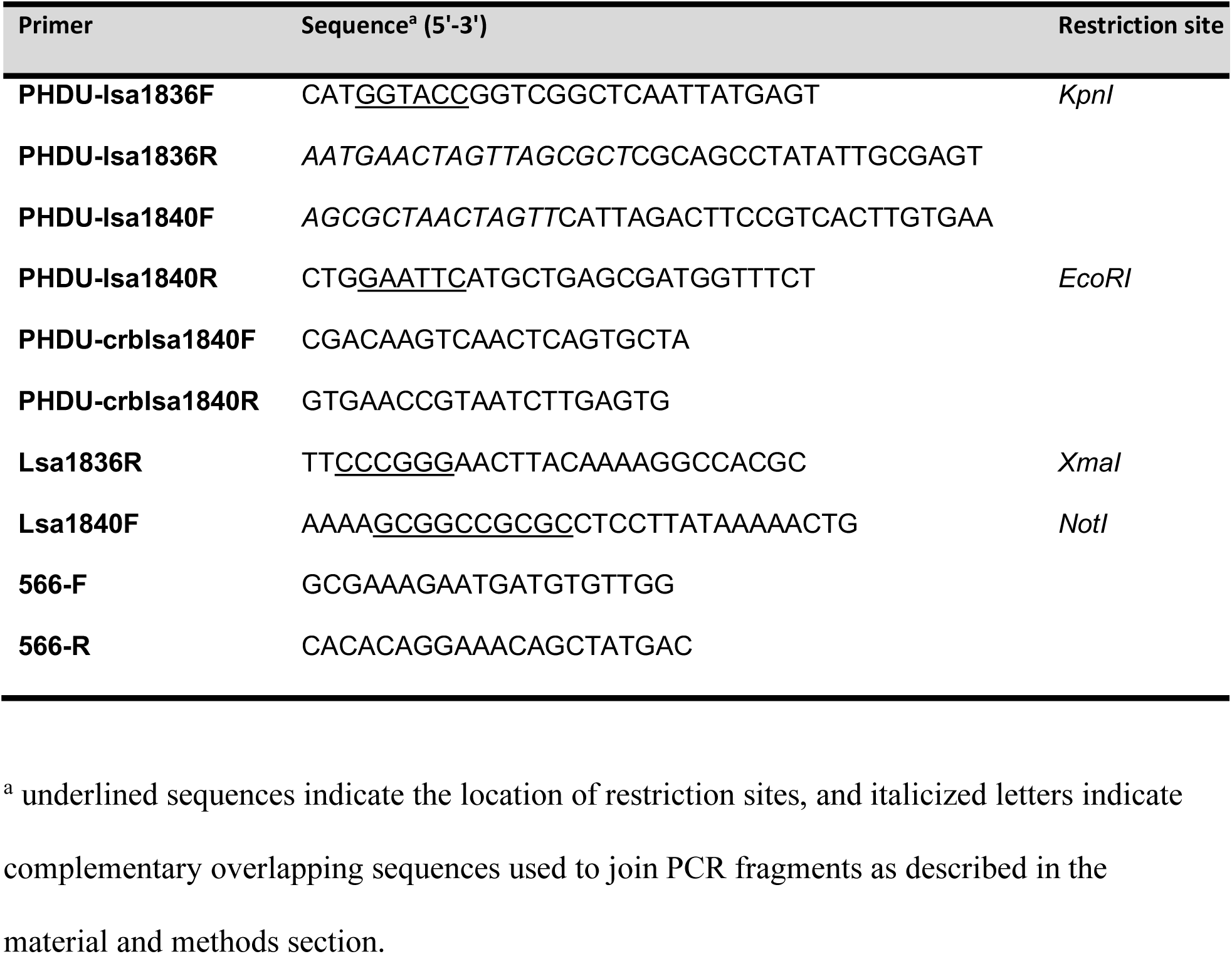
Oligonucleotides used in this study

### Bioinformatic analyses

Analyses were performed in the sequenced *L. sakei* 23K genome as described in (12). Each fasta sequence of every gene of the operon comprised between *lsa1836* and *lsa1840* was retrieved from UnitProtKB server at http://www.uniprot.org/uniprot, uploaded then analyzed using HHpred server (44) that detects structural homologues. For Lsa1839 and Lsa1840, that partly shares strong structural homology with Geranyl-geranyltransferase type-I (pdb id 5nsa, chain A) (51), and β domain of human haptocorrin (pdb id 4kki chain A) (52), intrinsic factor with cobalamin (pdb id 2pmv) (53) and transcobalamin (pdb id 2bb6 chainA) (54) respectively, homology modeling was performed using Modeler, version Mod9v18 (55). The heterodimer was then formed with respect to the functional and structural assembly of α and β domains of the native haptocorrin (52). Upon dimer formation, the best poses for heme within the groove, located at the interface of this heterodimer, were computed using Autodock4 tool (56). The protocol and grid box were previously validated with the redocking of cyanocobalamin within human haptocorrin (4kki) (42) and of cobalamin within bovine transcobalamin (2bb6). To compute the binding energy of every complex, the parameters of the cobalt present in the cobalamin and cyanocobalamin were added to the parameter data table, the iron parameters of the heme are already in the parameter data table. Then the docking poses were explored using the Lamarckian genetic algorithm. The poses of the ligands were subsequently analyzed with PyMOL of the Schrödinger suite (57). Comparative genomic analysis for conservation of gene synteny between meat-borne bacteria was carried out with the MicroScope Genome Annotation plateform, using the Genome Synteny graphical output and the PkGDB Synteny Statistics (58)

### Construction of plasmids and *L. sakei* mutant strains

All the primers and plasmids used in this study are listed in Table 2 and 3. The *lsa1836-1840* genes were inactivated by a 5118 bp deletion using double cross-over strategy. Upstream and downstream fragments were obtained using primers pairs PHDU-lsa1836F/PHDU-lsa1836R (731 bp) and PHDU-lsa1840F/PHDU-lsa1840R (742 bp) (Table 3). PCR fragments were joined by SOE using primers PHDU-lsa1836F/PHDU-lsa1840R and the resulting 1456 bp fragment was cloned between *EcoR*I and *KpnI*I sites in pRV300 yielding the pRV441 (Table 2). pRV441 was introduced in the *L. sakei* 23K and the *L. sakei* 23K Δ*lacLM* (RV2002) strains by electroporation as described previously (59). Selection was done on erythromycin sensitivity. Second cross-over erythromycin sensitive candidates were screened using primers PHDU-crblsa1840F and PHDU-crblsa1840R (Table 3). Deletion was then confirmed by sequencing the concerned region and the *lsa1836-1840* mutant strains were named RV4056 and RV4057 (Table 2).

To construct the RV2002 *hrtR-lac* and the RV4057 *hrtR-lac* strains, the pP_*hrt*_*hrtR-lac* (Table 2) was transformed by electroporation into the corresponding mother strains. For complementation, a pP*lsa1836-1840* plasmid (Table 2) was constructed as follows: a DNA fragment encompassing the promoter and the 5 genes of the *lsa1836-1840* operon was PCR amplified using the primers pair Lsa1836R/Lsa1840F (Table 3). The 5793 bp amplified fragment was cloned into plasmid pRV566 at *Xma*I and *Not*I sites. The construct was verified by sequencing the whole DNA insert using the 566-F and 566-R primers (Table 3) as well as internal primers. The pP*lsa1836-1840* was introduced into RV4056 bacteria by electroporation and transformed bacteria were selected for erythromycin resistance, yielding the RV4056c complemented mutant strain.

### β-galactosidase assay

Liquid cultures were usually grown in MCD into exponentially phase corresponding to a *A*_600_ equal to 0,5-0.8 and then incubated for 1 h at 30°C with hemin at the indicated concentration. β-Galactosidase (β-Gal) activity was assayed on bacteria permeabilized as described. β-Gal activity was quantified by luminescence in an Infinite M200 spectroluminometer (Tecan) using the β-Glo® assay system as recommended by manufacturer (Promega).

### Intracellular iron ^57^Fe determination

The various strains were grown in MCD to *A*_600_ = 0.5-0.7 at 30°C prior to addition or not of 0.1, 1, 5 or 40 µM ^57^Fe-labelled hemin (Frontier Scientific). Cells were then incubated at 30°C for an additional hour and overnight (19 hours). Cells were washed three times in H_2_O supplemented with 1mM EDTA. Cell pellets were desiccated and mineralized by successive incubations in 65% nitric acid solution at 130°C. ^57^Fe was quantified by Inductively Coupled Plasma Mass Spectroscopy (ICP-MS) (Agilent 7700X), Géosciences, University of Montpellier (France).

## Acknowledgments

This work, including Emilie Verplaetse post-doctoral grant, was funded from the French National Research Agency ANR-11-IDEX-0003-02; ‘ALIAS’ project. The authors would like to thank Véronique Martin for her help in setting up the cobalt parameter in the Autodock table parameter, Elise Abi-Khalil for the construction of the pLsa1836-1840, Delphine Lechardeur and Alexandra Gruss for the heme reporter plasmid and fruitful discussion and support.

## References

1. Neilands JB. 1981. Microbial Iron Compounds. Annu Rev Biochem 50:715–731.

2. Brooijmans R, Smit B, Santos F, van Riel J, de Vos WM, Hugenholtz J. 2009. Heme and menaquinone induced electron transport in lactic acid bacteria. Microb Cell Factories 8:28.

3. Pandey A, Bringel F, Meyer J-M. 1994. Iron requirement and search for siderophores in lactic acid bacteria. Appl Microbiol Biotechnol 40:735–739.

4. Champomier MC, Montel MC, Grimont F, Grimont PA. 1987. Genomic identification of meat *Lactobacilli* as *Lactobacillus sake*. Ann Inst Pasteur Microbiol 138:751–758.

5. Bredholt S, Nesbakken T, Holck A. 1999. Protective cultures inhibit growth of *Listeria monocytogenes* and *Escherichia coli* O157:H7 in cooked, sliced, vacuum- and gas-packaged meat. Int J Food Microbiol 53:43–52.

6. Leroy F, Lievens K, De Vuyst L. 2005. Modeling Bacteriocin Resistance and Inactivation of *Listeria innocua* LMG 13568 by *Lactobacillus sakei* CTC 494 under Sausage Fermentation Conditions. Appl Environ Microbiol 71:7567–7570.

7. Vermeiren L, Devlieghere F, Debevere J. 2004. Evaluation of meat born lactic acid bacteria as protective cultures for the biopreservation of cooked meat products. Int J Food Microbiol 96:149–164.

8. Chaillou S, Christieans S, Rivollier M, Lucquin I, Champomier-Vergès MC, Zagorec L. 2014. Quantification and efficiency of Lactobacillus sakei strain mixtures used as protective cultures in ground beef. Meat Sci 97:332–338.

9. Devlieghere F, Francois K, Vereecken KM, Geeraerd AH, Van Impe JF, Debevere J. 2004. Effect of chemicals on the microbial evolution in foods. J Food Prot 67:1977–1990.

10. Lombardi-Boccia G, Martinez-Dominguez B, Aguzzi A. 2002. Total Heme and Non-heme Iron in Raw and Cooked Meats. J Food Sci 67:1738–1741.

11. Hertel C, Schmidt G, Fischer M, Oellers K, Hammes WP. 1998. Oxygen-Dependent Regulation of the Expression of the Catalase Gene katA of Lactobacillus sakei LTH677. Appl Environ Microbiol 64:1359–1365.

12. Chaillou S, Champomier-Vergès M-C, Cornet M, Crutz-Le Coq A-M, Dudez A-M, Martin V, Beaufils S, Darbon-Rongère E, Bossy R, Loux V, Zagorec M. 2005. The complete genome sequence of the meat-borne lactic acid bacterium *Lactobacillus sakei* 23K. Nat Biotechnol 23:1527–1533.

13. Duhutrel P, Bordat C, Wu T-D, Zagorec M, Guerquin-Kern J-L, Champomier-Verges M-C. 2010. Iron Sources Used by the Nonpathogenic Lactic Acid Bacterium *Lactobacillus sakei* as Revealed by Electron Energy Loss Spectroscopy and Secondary-Ion Mass Spectrometry. Appl Environ Microbiol 76:560–565.

14. Huang W, Wilks A. 2017. Extracellular Heme Uptake and the Challenge of Bacterial Cell Membranes. Annu Rev Biochem 86:799–823.

15. Gruss A, Borezée-Durant E, Lechardeur D. 2012. Chapter Three - Environmental Heme Utilization by Heme-Auxotrophic Bacteria, p. 69–124. In Robert K. Poole (ed.), Advances in Microbial Physiology. Academic Press.

16. Choby JE, Skaar EP. 2016. Heme Synthesis and Acquisition in Bacterial Pathogens. J Mol Biol 428:3408–3428.

17. Anzaldi LL, Skaar EP. 2010. Overcoming the Heme Paradox: Heme Toxicity and Tolerance in Bacterial Pathogens. Infect Immun 78:4977–4989.

18. Reniere ML, Torres VJ, Skaar EP. 2007. Intracellular metalloporphyrin metabolism in *Staphylococcus aureus*. BioMetals 20:333–345.

19. Honsa ES, Maresso AW, Highlander SK. 2014. Molecular and Evolutionary Analysis of NEAr-Iron Transporter (NEAT) Domains. PLoS ONE 9:e104794.

20. Bates CS, Montanez GE, Woods CR, Vincent RM, Eichenbaum Z. 2003. Identification and Characterization of a *Streptococcus pyogenes* Operon Involved in Binding of Hemoproteins and Acquisition of Iron. Infect Immun 71:1042–1055.

21. Meehan M, Burke FM, Macken S, Owen P. 2010. Characterization of the haem-uptake system of the equine pathogen *Streptococcus equi subsp. equi*. Microbiology 156:1824–1835.

22. Lei B, Smoot LM, Menning HM, Voyich JM, Kala SV, Deleo FR, Reid SD, Musser JM. 2002. Identification and Characterization of a Novel Heme-Associated Cell Surface Protein Made by *Streptococcus pyogenes*. Infect Immun 70:4494–4500.

23. Ouattara M, Bentley Cunha E, Li X, Huang Y-S, Dixon D, Eichenbaum Z. 2010. Shr of group A *streptococcus* is a new type of composite NEAT protein involved in sequestering haem from methaemoglobin: Haem uptake and reduction by Shr. Mol Microbiol 78:739–756.

24. Lechardeur D, Cesselin B, Liebl U, Vos MH, Fernandez A, Brun C, Gruss A, Gaudu P. 2012. Discovery of Intracellular Heme-binding Protein HrtR, Which Controls Heme Efflux by the Conserved HrtB-HrtA Transporter in *Lactococcus lactis*. J Biol Chem 287:4752–4758.

25. Joubert L, Derré-Bobillot A, Gaudu P, Gruss A, Lechardeur D. 2014. HrtBA and menaquinones control haem homeostasis in *Lactococcus lactis*: Membrane and intracellular haem control in *Lactococcus lactis*. Mol Microbiol 93:823–833.

26. Rempel S, Stanek WK, Slotboom DJ. 2019. ECF-Type ATP-Binding Cassette Transporters. Annu Rev Biochem 88:551–576.

27. Finkenwirth F, Eitinger T. 2019. ECF-type ABC transporters for uptake of vitamins and transition metal ions into prokaryotic cells. Res Microbiol.

28. Wang T, Fu G, Pan X, Wu J, Gong X, Wang J, Shi Y. 2013. Structure of a bacterial energy-coupling factor transporter. Nature 497:272–276.

29. Bao Z, Qi X, Hong S, Xu K, He F, Zhang M, Chen J, Chao D, Zhao W, Li D, Wang J, Zhang P. 2017. Structure and mechanism of a group-I cobalt energy coupling factor transporter. Cell Res 27:675–687.

30. Zhang M, Bao Z, Zhao Q, Guo H, Xu K, Wang C, Zhang P. 2014. Structure of a pantothenate transporter and implications for ECF module sharing and energy coupling of group II ECF transporters. Proc Natl Acad Sci U S A 111:18560–18565.

31. Xu K, Zhang M, Zhao Q, Yu F, Guo H, Wang C, He F, Ding J, Zhang P. 2013. Crystal structure of a folate energy-coupling factor transporter from Lactobacillus brevis. Nature 497:268–271.

32. Rodionov DA, Hebbeln P, Eudes A, ter Beek J, Rodionova IA, Erkens GB, Slotboom DJ, Gelfand MS, Osterman AL, Hanson AD, Eitinger T. 2009. A novel class of modular transporters for vitamins in prokaryotes. J Bacteriol 191:42–51.

33. Henderson GB, Zevely EM, Huennekens FM. 1979. Mechanism of folate transport in Lactobacillus casei: evidence for a component shared with the thiamine and biotin transport systems. J Bacteriol 137:1308–1314.

34. Burgess CM, Slotboom DJ, Geertsma ER, Duurkens RH, Poolman B, van Sinderen D. 2006. The riboflavin transporter RibU in Lactococcus lactis: molecular characterization of gene expression and the transport mechanism. J Bacteriol 188:2752–2760.

35. Turner MS, Tan YP, Giffard PM. 2007. Inactivation of an Iron Transporter in *Lactococcus lactis* Results in Resistance to Tellurite and Oxidative Stress. Appl Environ Microbiol 73:6144–6149.

36. Li L, Chen OS, Ward DM, Kaplan J. 2001. CCC1 Is a Transporter That Mediates Vacuolar Iron Storage in Yeast. J Biol Chem 276:29515–29519.

37. Fernandez A, Lechardeur D, Derré-Bobillot A, Couvé E, Gaudu P, Gruss A. 2010. Two Coregulated Efflux Transporters Modulate Intracellular Heme and Protoporphyrin IX Availability in *Streptococcus agalactiae*. PLoS Pathog 6:e1000860.

38. von Heijne G. 1992. Membrane protein structure prediction. Hydrophobicity analysis and the positive-inside rule. J Mol Biol 225:487–494.

39. Jin B, Newton SMC, Shao Y, Jiang X, Charbit A, Klebba PE. 2006. Iron acquisition systems for ferric hydroxamates, haemin and haemoglobin in *Listeria monocytogenes*. Mol Microbiol 59:1185–1198.

40. Abi-Khalil E, Segond D, Terpstra T, Andre-Leroux G, Kallassy M, Lereclus D, Bou-Abdallah F, Nielsen-Leroux C. 2015. Heme interplay between IlsA and IsdC: Two structurally different surface proteins from *Bacillus cereus*. Biochim Biophys Acta 1850:1930–1941.

41. Maresso AW, Chapa TJ, Schneewind O. 2006. Surface Protein IsdC and Sortase B Are Required for Heme-Iron Scavenging of *Bacillus anthracis*. J Bacteriol 188:8145–8152.

42. Mazmanian SK, Skaar EP, Gaspar AH, Humayun M, Gornicki P, Jelenska J, Joachmiak A, Missiakas DM, Schneewind O. 2003. Passage of heme-iron across the envelope of *Staphylococcus aureus*. Science 299:906–909.

43. Mazmanian SK, Ton-That H, Su K, Schneewind O. 2002. An iron-regulated sortase anchors a class of surface protein during *Staphylococcus aureus* pathogenesis. Proc Natl Acad Sci 99:2293–2298.

44. Söding J, Biegert A, Lupas AN. 2005. The HHpred interactive server for protein homology detection and structure prediction. Nucleic Acids Res 33:W244–248.

45. Szklarczyk D, Gable AL, Lyon D, Junge A, Wyder S, Huerta-Cepas J, Simonovic M, Doncheva NT, Morris JH, Bork P, Jensen LJ, Mering C von. 2019. STRING v11: protein– protein association networks with increased coverage, supporting functional discovery in genome-wide experimental datasets. Nucleic Acids Res 47:D607–D613.

46. Roberts JN, Singh R, Grigg JC, Murphy MEP, Bugg TDH, Eltis LD. 2011. Characterization of Dye-Decolorizing Peroxidases from *Rhodococcus jostii* RHA1. Biochemistry 50:5108–5119.

47. Singh R, Grigg JC, Armstrong Z, Murphy MEP, Eltis LD. 2012. Distal Heme Pocket Residues of B-type Dye-decolorizing Peroxidase: ARGININE BUT NOT ASPARTATE IS ESSENTIAL FOR PEROXIDASE ACTIVITY. J Biol Chem 287:10623–10630.

48. Lauret R, Morel-Deville F, Berthier F, Champomier-Verges M, Postma P, Ehrlich SD, Zagorec M. 1996. Carbohydrate utilization in *Lactobacillus sake*. Appl Environ Microbiol 62:1922–1927.

49. Gaudu P, Vido K, Cesselin B, Kulakauskas S, Tremblay J, Rezaiki L, Lamberret G, Sourice S, Duwat P, Gruss A. 2002. Respiration capacity and consequences in *Lactococcus lactis*. Antonie Van Leeuwenhoek 82:263–269.

50. Lechardeur D, Cesselin B, Fernandez A, Lamberet G, Garrigues C, Pedersen M, Gaudu P, Gruss A. 2011. Using heme as an energy boost for lactic acid bacteria. Curr Opin Biotechnol 22:143–149.

51. Bloch JS, Ruetz M, Kräutler B, Locher KP. 2017. Structure of the human transcobalamin beta domain in four distinct states. PLOS ONE 12:e0184932.

52. Furger E, Frei DC, Schibli R, Fischer E, Prota AE. 2013. Structural Basis for Universal Corrinoid Recognition by the Cobalamin Transport Protein Haptocorrin. J Biol Chem 288:25466–25476.

53. Mathews FS, Gordon MM, Chen Z, Rajashankar KR, Ealick SE, Alpers DH, Sukumar N. 2007. Crystal structure of human intrinsic factor: Cobalamin complex at 2.6-A resolution. Proc Natl Acad Sci 104:17311–17316.

54. Wuerges J, Garau G, Geremia S, Fedosov SN, Petersen TE, Randaccio L. 2006. Structural basis for mammalian vitamin B12 transport by transcobalamin. Proc Natl Acad Sci 103:4386–4391.

55. Webb B, Sali A. 2016. Comparative Protein Structure Modeling Using MODELLER. Curr Protoc Bioinforma 54:5.6.1-5.6.37.

56. Morris GM, Goodsell DS, Halliday RS, Huey R, Hart WE, Belew RK, Olson AJ. 1998. Automated docking using a Lamarckian genetic algorithm and an empirical binding free energy function. J Comput Chem 19:1639–1662.

57. The PyMOL Molecular Graphics System, Version 2.0. Schrödinger, LLC.

58. Vallenet D, Belda E, Calteau A, Cruveiller S, Engelen S, Lajus A, Le Fèvre F, Longin C, Mornico D, Roche D, Rouy Z, Salvignol G, Scarpelli C, Thil Smith AA, Weiman M, Médigue C. 2013. MicroScope—an integrated microbial resource for the curation and comparative analysis of genomic and metabolic data. Nucleic Acids Res 41:D636–D647.

59. Berthier F, Zagorec M, Champomier-Verges M, Ehrlich SD, Morel-Deville F. 1996. Efficient transformation of Lactobacillus sake by electroporation. Microbiology 142:1273–1279.

60. Stentz R, Loizel C, Malleret C, Zagorec M. 2000. Development of Genetic Tools for *Lactobacillus sakei*: Disruption of the β-Galactosidase Gene and Use of *lacZ* as a Reporter Gene To Study Regulation of the Putative Copper ATPase, AtkB. Appl Environ Microbiol 66:4272–4278.

61. Leloup L, Ehrlich SD, Zagorec M, Morel-Deville F. 1997. Single-crossover integration in the *Lactobacillus sake* chromosome and insertional inactivation of the *ptsI* and *lacL* genes. Appl Environ Microbiol 63:2117–2123.

62. Alpert C-A, Crutz-Le Coq A-M, Malleret C, Zagorec M. 2003. Characterization of a Theta-Type Plasmid from *Lactobacillus sakei*: a Potential Basis for Low-Copy-Number Vectors in *Lactobacilli*. Appl Environ Microbiol 69:5574–5584.

