## supplemental tables S1, S2, Fig SI S2 for "Heme uptake in *Lactobacillus sakei* evidenced by a new ECF-like transport system"

Running title: heme transport in *Lactobacillus sakei*

Key-words: iron, lactic acid bacteria, ABC-transporter

**Results**

In order to investigate the link between the presence of heme in the growth medium and survival in *L. sakei*, we grew bacterial cells under various heme concentrations and determined: i) cell survival and ii) iron accumulation in cell cytoplasm. As shown in Figure S1, increasing heme concentration leads to an increased cell survival during the stationary growth phase. This is correlated with an increased iron accumulation into the wild-type (WT) 23K cells, as revealed by EELS analysis on cells grown in 5 µM and 40 µM heme (Figure S2). This analysis reveals that there is more intracellular iron accumulated when cells are grown with 40 µM heme compared to 5 µM. These differences in iron cellular contents are correlated with bacterial survival. Increasing heme concentration over 40 µM and up to 120 µM did not result in enhanced survival which becomes maximal for 40 µM heme. Nonetheless, these elevated heme concentrations did not appear to be toxic for *L. sakei* cells since no more morbidity was observed between 40 and 120 µM heme (Figure S1).

**The LSA1194-95 is an actor of heme uptake in *Lactobacillus sakei in vitro.***

The involvement of the *lsa1194-1195* gene product in heme transport across the membrane was also investigated using the intracellular heme sensor. A *lsa1194-1195* mutant was constructed in the Δ*lacLM* RV2002 genetic background yielding the RV4070 strain (Table S1). The $pP_{hrt}$ *hrtR-lac* was then introduced in the RV4070 strain, yielding the RV4057 *hrtR-lac* strain (Table S1). β-Galactosidase (β-Gal) activity of the RV4070 *hrtR-lac* strain grown in MCD in the presence of 0.5, 1 and 5 µM hemin was determined and compared to that of the RV2002 *hrtR-lac* used as control (Fig. S3A). Relative β-Gal activity of the RV4070 *hrtR-lac* strain showed a slight increase as compared to the RV2002 strain at 0.5 µM heme however with great variability between samples as can be seen with big error bars. A two-fold reduction was measured at 1 µM heme and still a 35% reduced activity was shown at higher hemin concentration. This indicates that the intracellular abundance of heme is reduced in the RV4057 bacterial cells at 1 and 5 µM heme while it is similar to the RV2002 strain at lower heme concentrations. To quantify the absolute amount of heme incorporated by bacteria lacking this putative transporter, the *lsa1194-1195* mutant was constructed in the WT *L. sakei* 23K genetic background yielding the RV4069 strain (Table S1). 23K and RV4069 cells were incubated in the MCD in the absence of in the presence of 1, 5 or 40 µM of $^{57}$Fe-hemin. ICP-MS quantification indicated that the $^{57}$Fe content of the two strains was similar at 1 µM, 5 µM and 40 µM $^{57}$Fe-hemin (Fig. S3B).

**The *lsa1194-1195* mutant strain is affected in its survival in the presence of heme.**

Like other lactic acid bacteria, *L. sakei* does not require heme to grow. However, its survival was highly enhanced during stationary phase when cells were grown in the presence of heminic compounds (hematin, myoglobin, and hemoglobin), with cells being able to survive up to 7 days (13). The increase of heme concentration in the growth medium being correlated with an increase in the incorporation of heme and a prolong viability of cells, one may expect that the *lsa1836-1840,* or *lsa1194-95,* mutant strains are affected in their survival. Viability of the RV4057 and the RV4069 grown in the MCD in the absence and in the presence of 40 µM heme was assessed. Survival of *L. sakei* was assessed in the presence of both hemin or hematin molecules and gave a similar effect on *L. sakei* cell viability. This heme concentration appeared to promote the highest survival rate of bacteria (Fig. S1). Despite the defect in heme incorporation (Fig. 3), we did not notice a survival defect of the RV4056 strain as cells still display a wild-type physiological response (Fig. S4A). As for the RV4069, a small but reproducible decrease in the survival population was observed (Fig. S4B).

### Material and methods

**Bacterial strains and growth conditions:** *L. sakei* 23K (Chaillou et al., 2005) was propagated on MRS medium at 30°C (De Man et al., 1960). For physiological studies the chemically defined medium MCD (Lauret et al., 1996) supplemented with 0.5% (wt/vol) glucose was used. MCD medium contains no iron sources but contains possible traces of iron coming from various components or distilled water. Incubation was performed at 30°C with stirring at 70 rpm. *E. coli* strains were grown at 37°C under aerobiosis in Luria Bertani broth (LB). *L. sakei* cell growth and viability were followed by measuring the optical density at 600 nm (OD600) on a visible spectrophotometer (Secoman) and by the determination of the number of colonies (cfu.ml-1) after plating serial dilutions of samples on MRS agar. Plates were incubated under aerobiosis at 30°C for 30 h. All the measurements were performed in at least three independent assays. When needed, media were supplemented with freshly prepared filtered solution to obtain final concentrations of 1, 5, 20, 40, 80 or 120 µM hematin or hemin (Sigma- Aldrich). Ampicillin (100 $\mu g.ml^{-1}$) was used for the selection of *E. coli* strains. Erythromycin (5 $\mu g.ml^{-1}$) was used for the selection of *L. sakei* strains. Serial dilutions were done in Dilution Medium [Bacto Beef extract ($5g.l^{-1}$, BD) and universal peptone M66 (15 $g.L^{-1}$, Merck)

**Intra cellular iron mapping:** It was determined by EELS analysis on cells grown 8h in MCD medium (Lauret et al., 1996) supplemented with hematin (5 or 40 µM) or not, as previously described (Duhutrel et al., 2010).

**Construction of the *L. sakei lsa1194-1195* mutant strains.**

*lsa1194-1195* genes were inactivated by a 1344 bp deletion using double cross-over strategy. Upstream and downstream fragments were obtained using primers pairs PHDU-lsa1194F and PHDU-lsa1194R (939 bp) and PHDU-lsa1195F and PHDU-lsa1195R (891 bp) (Table S2), respectively. They were joigned using primers PHDU-lsa1194F and PHDU-lsa1195R leading to a 1830 bp fragment encompassing a sequence complementarity of about 21 bp between upstream and downstream fragments. The amplified fragment was cloned into plasmid pRV300 at *Kpn*I and *Pst*I sites to form plasmid pRV446. *E. coli* DH5α was used for its multiplication. This plasmid was introduced into the *L. sakei* 23K and RV2002 strains (Table 1) by electroporation as described previously (Berthier *et al.*, 1996). Selection was done on erythromycin sensitivity. Second cross-over erythromycin sensitive candidates were screened using primers PHDU-lsa1194F and PHDU-crblsa1195R (Table S2) to detect *lsa1194-1195* deletion which led to a 1208 bp fragment compare to 2528 bp for WT allele. Deletion was then confirmed by sequencing the concerned region and the *lsa1194-1195* mutant strains were respectively named RV4069 and RV4070 (Table S1).

To construct the RV4070 *hrtR-lac* strain, the $pP_{hrt}$*hrtR-lac* (Table 1) was transformed by electroporation into the corresponding mother strain (Table S1).

Table S1: Strains and plasmids used in this study

| Strains or plasmids | Characteristics | References |
|---|---|---|
| **Strains** | | |
| **RV4069** | 23K derivative, $\Delta$*lsa1194-1195* | This study |
| **RV4070** | RV2002 $\Delta$*lsa1194-1195* | This study |
| **RV4070 hrtR-lac** | RV4070 carrying the $pP_{hrt}$ *hrtR-lac*, $ery^{R}$ | This study |
| **Plasmids** | | |
| **pRV446** | pRV300 derivative, exchange cassette for *lsa1194-1195* | This study |

Table S2: Primers used in this study

| Primer | Sequence[a] (5'-3') | Restriction site |
|---|---|---|
| **PHDU-lsa1194F** | CGATGGTACCGCAAATTCACCAATCTTATAATG | *Kpn*I |
| **PHDU-lsa1194R** | *GATTGACTAGTTGCAGCG*TACGCGCATGACGTTAATTTTTTG | |
| **PHDU-lsa1195F** | *CGTACGCTGCAACTAGTC*AATCCACCATGGCTATTCACTAC | |
| **PHDU-lsa1195R** | ATGCCTGCAGGCCACCAGTCATTCTAAGT | *Pst*I |
| **PHDU-crblsa1195R** | TAGGAATTGCGCATATCAGT | |

[a] underlined sequences indicate the location of restriction sites, and italicized letters indicate complementary overlapping sequences used to join PCR fragments as described in the material and methods section.

Figure S1: Effects of heme concentration on long term survival of *L. sakei* cells. Heme concentrations: 1 µM (Full rhombus), 5 µM (Full squares), 40 µM (Open squares, dotted line), 80 µM (Full circles) and 120 µM (Open circles).

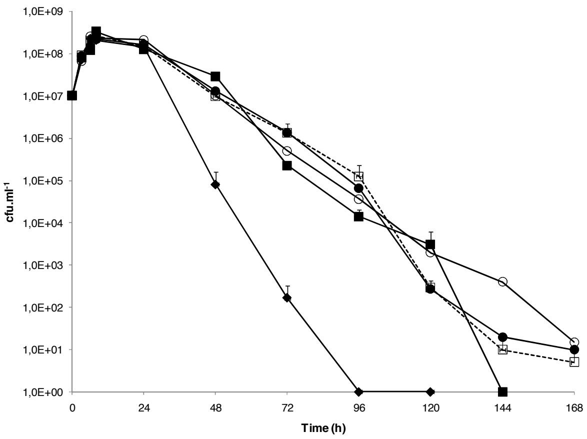

Figure S2: Iron mapping of *L. sakei* cells cultivated in MCD medium supplemented with heme A) 40 µM and B) 5 µM. ESI images obtained using EFTEM (cell scale). A filtered image at 250 eV (leading to inverted contrast) allows the morphology to be observed. The iron map, in red, is superimposed on the 250 eV image.

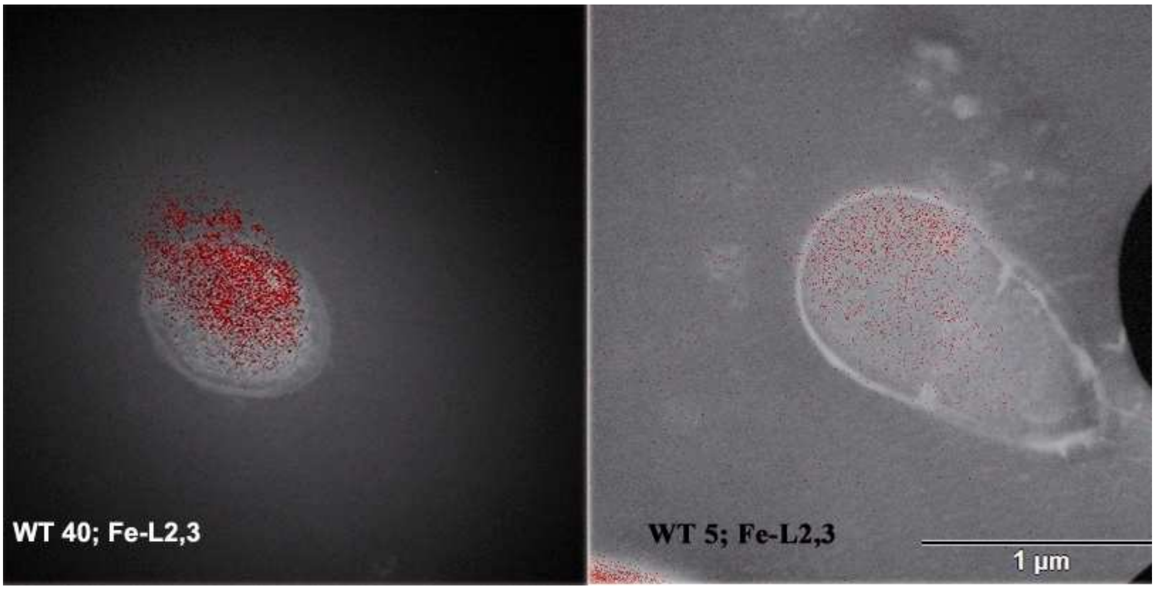

Figure S3: A, *In vivo* detection of intracellular heme content of the RV2002 and *lsa1194-1195* (RV4070) mutant strains. Strains carrying the p$P_{hrtR}$*hrtR-lac* were grown in hemin and β-Gal activity was quantified by luminescence (see "Material and methods"). For each experiment, values of luminescence obtained with no added hemin are subtracted and β-Gal activity of strains was expressed as the percentage to the RV2002 strain for each hemin concentration. Mean values are shown (n=3). Error bars represent the standard deviation. B, Quantification of the $^{57}$Fe content of the wild-type (23K) and the *lsa1194-1195* mutant (RV4069) strains grown in the absence and presence of indicated $^{57}$Fe-hemin concentrations. Results represent the mean and range from at least two independent experiments.

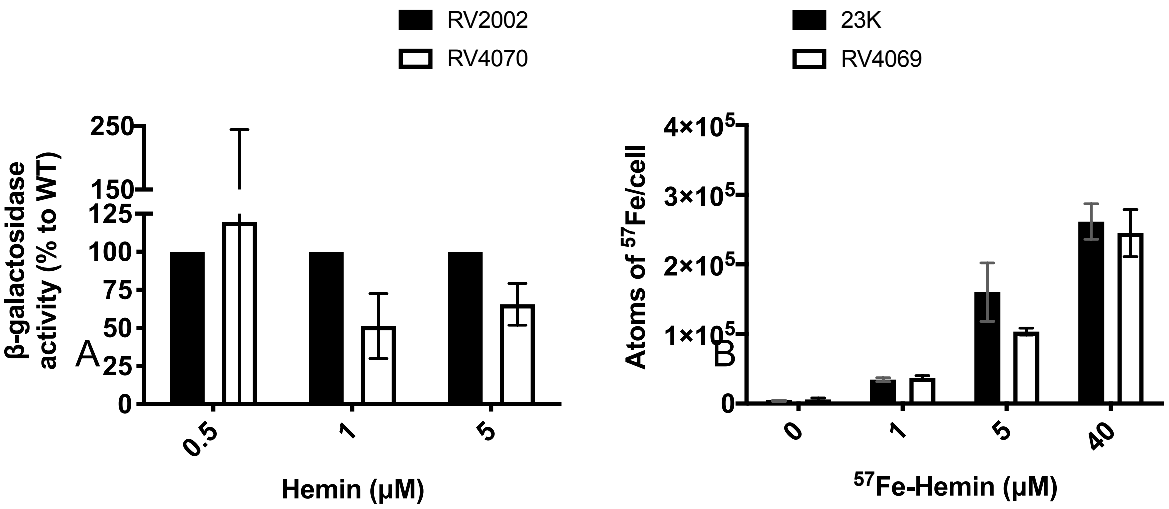

Figure S4: Survival of the *lsa1194-1195* mutant (RV4069) strain is reduced in the presence of heme. *lsa1836-1840* mutant (RV4056) (A) and lsa*1194-1195* mutant (RV4069) (B) viable cells grown in the presence of 40 µM hematin or hemin were enumerated every 24 hours over a period of 96 h. As a control, viable count of Ls 23K cells grown in the same growth conditions was performed.

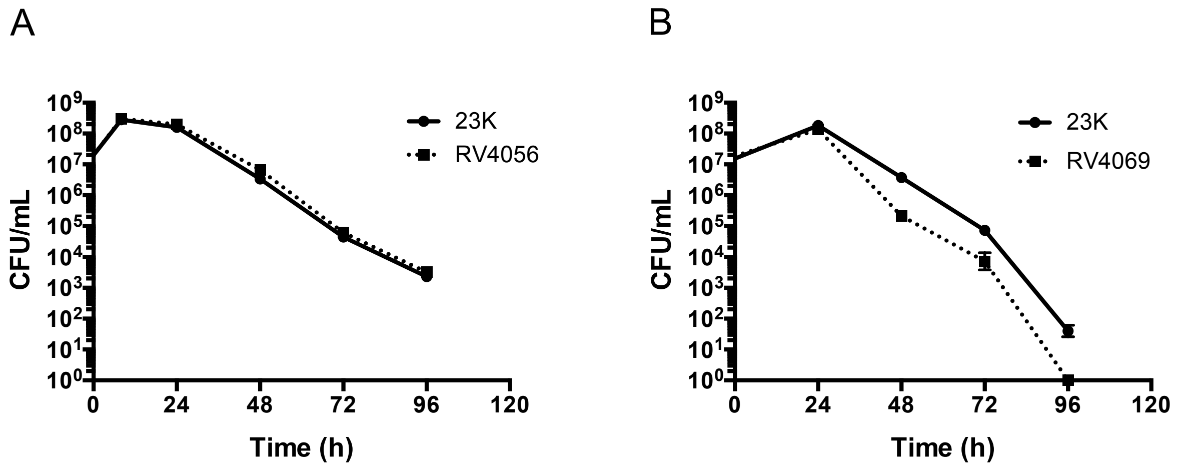
